# KSHV 3.0: A State-of-the-Art Annotation of the Kaposi’s Sarcoma-Associated Herpesvirus Transcriptome Using Cross-Platform Sequencing

**DOI:** 10.1101/2023.09.21.558842

**Authors:** István Prazsák, Dóra Tombácz, Ádám Fülöp, Gábor Torma, Gábor Gulyás, Ákos Dörmő, Balázs Kakuk, Lauren McKenzie Spires, Zsolt Toth, Zsolt Boldogkői

**Author notes:** Shared senior/corresponding authors. The first three authors contributed equally to this work.

## Abstract

Kaposi’s sarcoma-associated herpesvirus (KSHV) is a large, oncogenic DNA virus belonging to the gammaherpesvirus subfamily. KSHV has been extensively studied with various high-throughput RNA-sequencing approaches to map the transcription start and end sites, the splice junctions, and the translation initiation sites. Despite these efforts, the comprehensive annotation of the viral transcriptome remains incomplete. In the present study, we generated a long-read sequencing dataset of the lytic and latent KSHV transcriptome using native RNA and direct cDNA sequencing methods. This was supplemented with CAGE sequencing based on a short-read platform. We also utilized datasets from previous publications for our analysis. As a result of this combined approach, we have identified a number of novel viral transcripts and RNA isoforms and have either corroborated or improved the annotation of previously identified viral RNA molecules, thereby notably enhancing our comprehension of the transcriptomic architecture of the KSHV genome. We also evaluated the coding capability of transcripts previously thought to be non-coding, by integrating our data on the viral transcripts with translatomic information from other publications.

**IMPORTANCE:** Deciphering the viral transcriptome of KSHV is of great importance because we can gain insight into the molecular mechanism of viral replication and pathogenesis, which can help develop potential targets for antiviral interventions. Specifically, the identification of substantial transcriptional overlaps by this work suggests the existence of a genome-wide interference between transcriptional machineries. This finding indicates the presence of a novel regulatory layer, potentially controlling the expression of viral genes.

## INTRODUCTION

Infection with Kaposi’s sarcoma-associated herpesvirus (KSHV), a gamma-2 herpesvirus, is usually asymptomatic in healthy individuals, but it can lead to severe diseases in patients with compromised immune systems, such as those with AIDS (1). KSHV is the etiological agent of Kaposi’s sarcoma (2), which is characterized by the formation of aberrant blood vessels in the skin, mucous membranes, and internal organs (3). Beyond this, KSHV has also been associated with other forms of cancer, such as primary effusion lymphoma and multicentric Castleman’s disease (4).

KSHV is an enveloped, dsDNA virus with approximately 90 protein-coding genes in its genome. Extensive non-coding RNA (ncRNA) expression has also been detected in certain genomic regions (5). Its lifecycle comprises of a latent and lytic phase. Following infection of oral epithelial cells, KSHV eventually infects B lymphocytes, where it establishes latency, leading to a lifelong persistence in humans (6). KSHV can also establish latency in several cultured cell lines (7). During latency, KSHV produces four protein-coding genes, comprising of LANA, vCyclin, vFLIP and Kaposin encoded by ORF73, ORF72, ORF71 and by K12, respectively (8–10). Around 25 microRNAs (miRNAs) encoded by 12 precursor miRNAs (pre-miRNAs) (11–13) are also expressed during latency. LANA is responsible for maintaining the episomal viral DNA in the nucleus of infected cells (14, 15). KSHV miRNAs have been shown to target cellular mRNAs (11, 12, 16) and play a role in adjusting antiviral pathways (17) as well as influencing cellular growth. KSHV generates another type of ncRNA, known as circRNAs whose expression is induced during lytic infection (18, 19). The lytic cycle is initiated by the immediate early trans-regulator, known as RTA, encoded by ORF50 (20, 21). The let-7a miRNA/RBPJ signal is competitively modulated by RTA and LANA (22). RTA increases RBPJ levels by regulating let-7a, while LANA decreases RBPJ levels. During the lytic cycle, KSHV produces a significant amount of long non-coding RNA (lncRNA) called PAN spanning the K7 gene (23, 24).

KSHV transcripts have previously been examined using traditional techniques, including Northern blot, qPCR, and microarray analyses (25–28). Long-read sequencing (LRS) technologies have recently become important in viral transcriptomics (29–36). Native RNA sequencing is widely regarded as the gold standard in transcriptome research due to its ability to minimize the generation of spurious transcripts that can occur during library preparation and sequencing with other techniques. However, it is important to note that this technique also has certain limitations. For instance, transcript truncation might occur due to the digestion by RNase enzymes or during RNA preparation, which can lead to false identification of transcription start sites (TSSs). Using short-read sequencing (SRS) technology, high-resolution transcriptome maps of KSHV have been produced, leading to the discovery of novel bi-cistronic transcripts, splice variants, and gene expression dynamics during both latent and lytic infection phases (37–40). Aries and colleagues demonstrated that the long TSS isoforms can contain upstream ORFs (uORFs) encoding regulatory proteins (41). A recent RAMPAGE study (42) uncovered numerous previously unidentified TSSs and their associated cis-regulatory elements (43). To date, the number of KSHV introns has increased nearly tenfold, and numerous new splicing events have been detected for KSHV transcripts using ultra-deep SRS (44, 45).

Despite the popularity of SRS, this technique is limited in its ability to assemble full-length transcripts and accurately identify the isoform profiles. LRS approaches, such as Oxford Nanopore Technologies (ONT), overcome these limitations. They offer single-molecule sequencing with the capability to detect full-length transcripts and allow the accurate identification of transcript isoforms, including splice and length variants (29). Despite extensive high-throughput RNA-sequencing efforts to profile viral transcripts, the KSHV transcriptome is still incomplete. Here, we employed nanopore sequencing alongside other techniques to provide a comprehensive annotation of the KSHV transcripts in primary effusion lymphoma cells.

## RESULTS

### Methods for the analysis of the KSHV transcriptome

In this study, we employed a combined RNA sequencing approach to analyze the poly(A)^+^ fraction of both the lytic and latent KSHV transcriptomes in the transgenic PEL cell line iBCBL1-3xFLAG-RTA. Lytic reactivation was induced by adding doxycycline to the cells to express the 3xFLAG-RTA transgene, the inducer of the lytic cycle (**Figure 1**). The lytic viral transcriptome was analyzed at 24 hpi at which time point all lytic genes have been shown to be expressed. Our investigations incorporated direct cDNA- and native RNA-based library preparation techniques for nanopore sequencing (dcDNA-Seq and dRNA-Seq, respectively), along with Cap Analysis of Gene Expression sequencing (CAGE-Seq) which was carried out using an Illumina platform. The mapped reads were then subjected to transcript annotation applying the LoRTIA pipeline developed in our laboratory (46) (**Figure 2**). In this work, the LoRTIA program was employed for assessing the quality of sequencing adapters and poly(A) sequences, and filtering out false TSSs, TESs, and splice sites that could arise from RNA degradation, reverse transcription (RT), second strand synthesis, PCR amplification, or the sequencing reaction itself (46). The relevance of all eligible features was assessed against the Poisson or negative binomial distributions, and the p-value was adjusted using the Bonferroni correction method. Additional stringent filtering criteria were applied to enhance the confidence in the validity of the annotated LoRTIA transcripts (see below). In our pipeline, we also included a check for the potential presence of A-rich regions upstream of the transcriptional end sizes (TESs). Reads apparently originating from these regions were discarded as they may be indicative of false priming events. In order to further ensure that the annotated transcripts were not the result of possible internal priming events, we employed the talon_label_reads submodule of the TALON software package (47) for transcript annotation using the reads identified by the LoRTIA program. In this work, we also utilized datasets on KSHV from other publications (**Supplemental File 1**).

**Figure 1.**
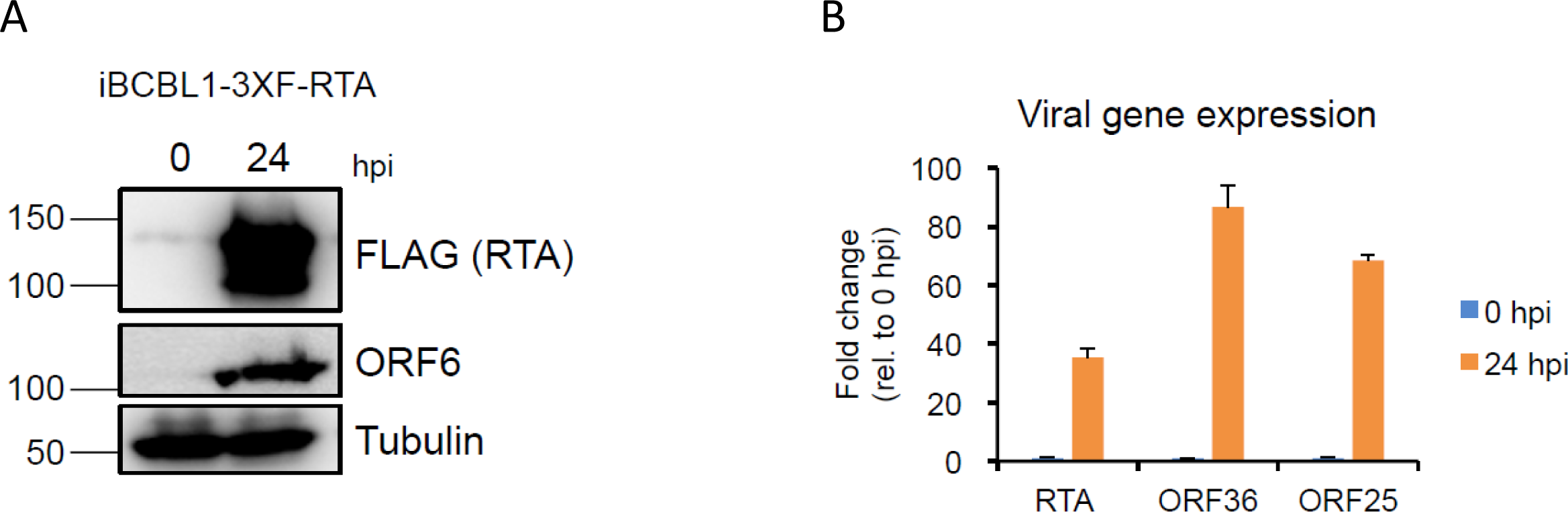
Induction of KSHV lytic infection. **(A)** Immunoblot analysis of the KSHV ORF6 protein and the housekeeping protein tubulin of the host cell at 0h and 24h post induction in iBCBL1-3xFLAG-RTA cells. **(B)** Quantitative RT-PCR analysis of immediate-early RTA, early ORF36 and late ORF25 viral mRNAs at 0h and 24h post induction.

**Figure 2.**
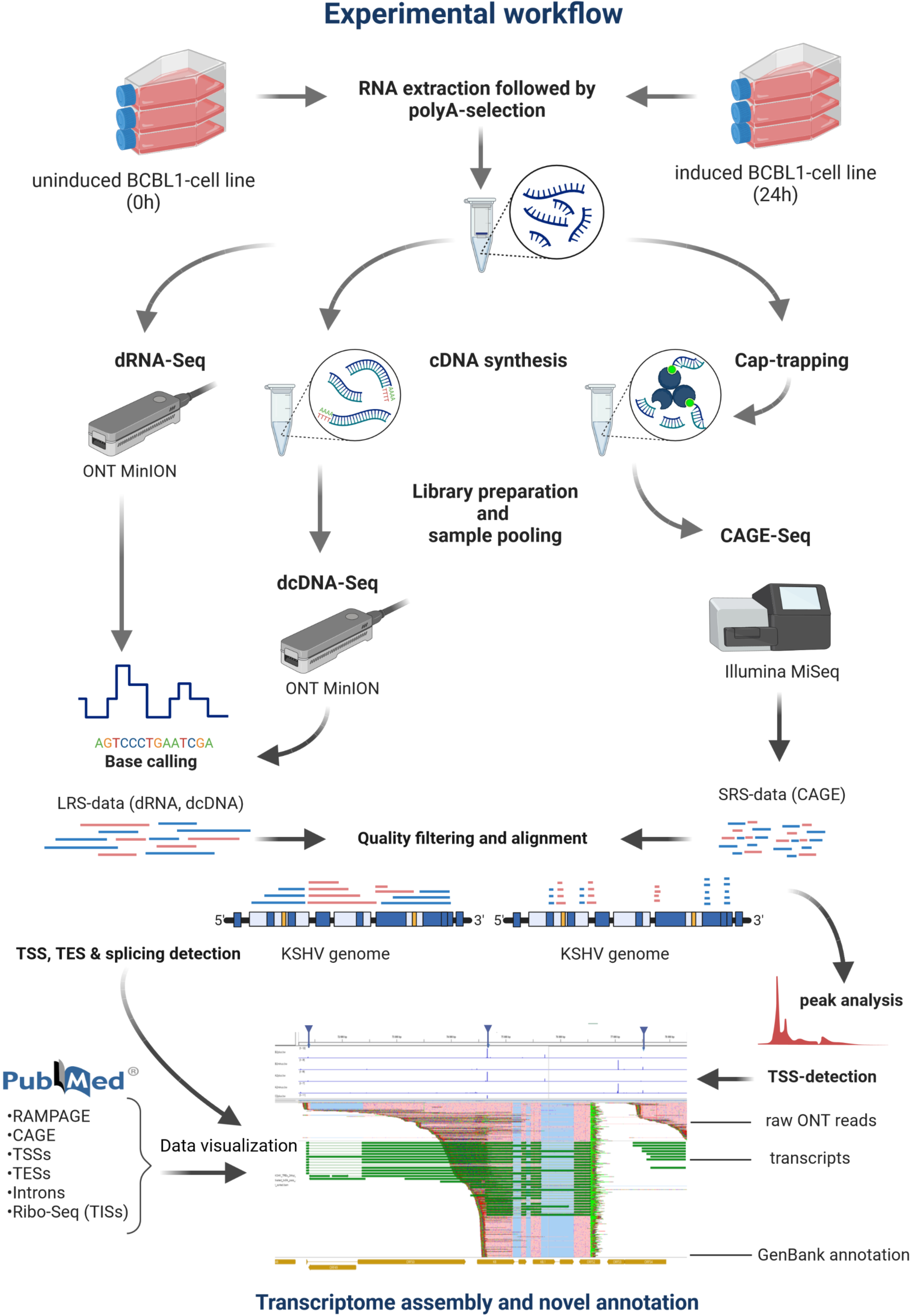
Workflow. This visualization details the methods used for generating novel sequencing data, encompassing steps such as DNA extraction, library preparation, RNA sequencing, and subsequent bioinformatics analyses. The flowchart illustrates each phase for clarity and order of execution.

### Identifying transcriptional start and end sites

The transcript ends were determined using LRS and CAGE-Seq methods, revealing an overrepresentation of PAN transcripts in the sample at 24 hpi (**Figure 3** and **Supplemental Figure 1**). Other transcripts, such as K7, K8, K8.1, ORF11, K2, and K4 mRNAs also exhibit a comparatively high count of transcript ends. We compared the obtained results with previously published data from RAMPAGE (43), 3’RACE (48), and CAGE-Seq (41) studies. As a result, we identified 557 positions as TSS from the dcDNA-Seq samples, with 408 of them satisfying our criteria for transcript annotation. Remarkably, 38% of these positions (211 out of 557) were independently confirmed by RAMPAGE and 5’ RACE with base pair precision (**Supplemental Table 1, Supplemental Figure 2**). In total, we identified 181 TESs using the LoRTIA software. Out of these, 146 were subsequently confirmed through dRNA sequencing. Additionally, we observed a poly(A) signal in 97 cases, positioned on average approximately 26.65 base pairs upstream of the TESs (**Supplemental Table 1)**. Among the 181 detected TES positions, 106 were deemed suitable as 3’ ends for transcript assembly. These TESs were also compared to already described transcript end positions. Notably, the TES positions exhibited significant overlap with the majority of pre-existing annotated TESs, encompassing 94% (64 out of 68) within a 10 base pair range (**Supplemental Table 1, Supplemental Figures 2**).

**Figure 3.**
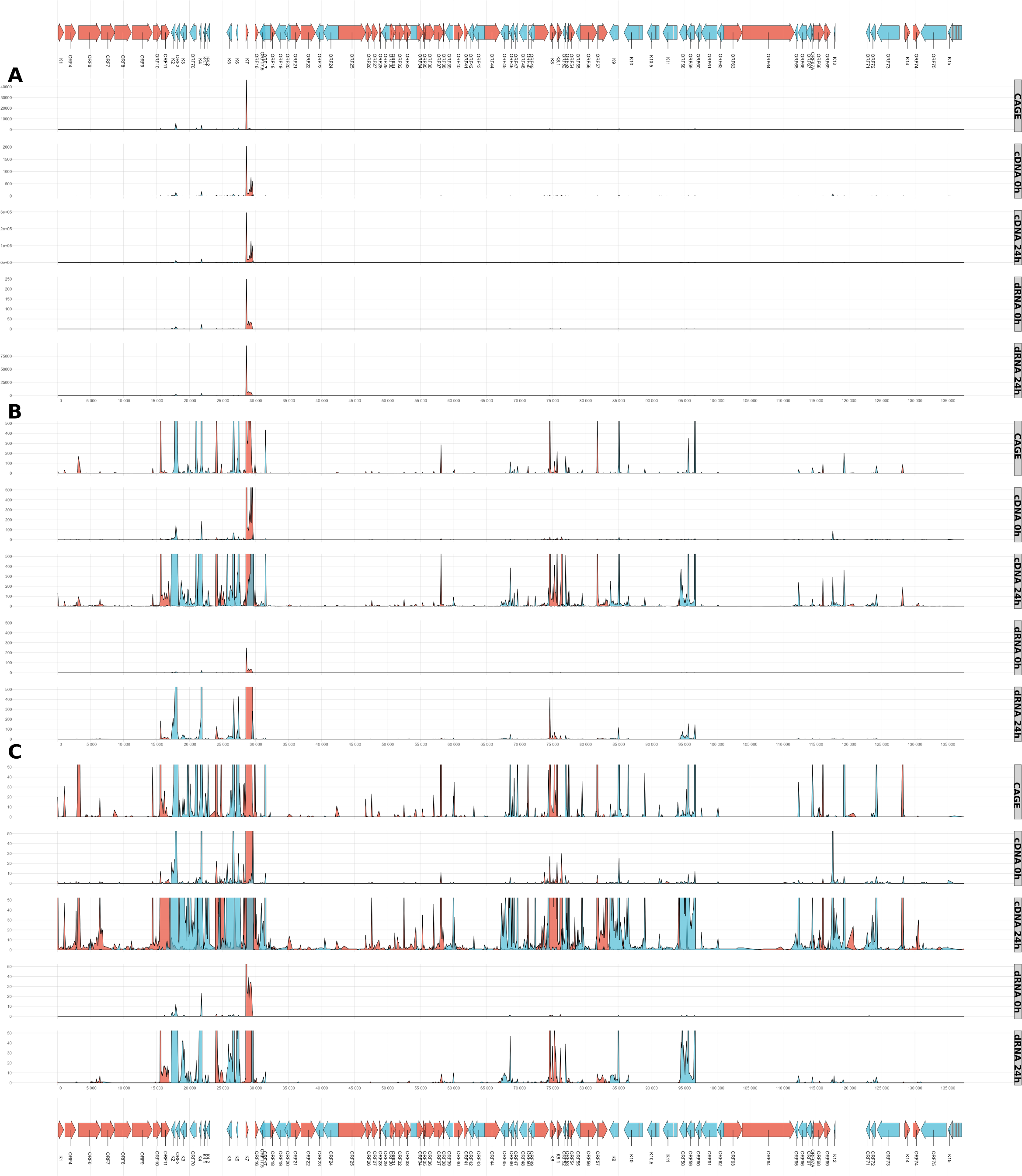
Transcriptional start sites of KSHV RNAs. TSS distributions derived from CAGE-Seq, dcDNA, and dRNA sequencing data. For dcDNA and dRNA methods, only the 24 hpi samples are illustrated. For each nucleotide the number of reads that started with their 5’ ends at the given position were counted. For the dcDNA-Seq, only reads with discernible directionality, identifiable by the presence of either 5’ or 3’ adapters, were included. For the dcDNA samples, data from all three replicates were merged. The TSS signal strength values were subsequently aggregated in 100-nt segments to present the distributions of TSSs. The y-axis limit was set to auto scale in (A) to 500 reads (B), and 50 reads (C). Gene (represented by arrows) and TSS distributions are differentiated by color: the positive strand is marked in red, while the negative strand is in blue.

The application of advanced 5’-end sequencing technologies revealed that most, if not all, promoters use tightly clustered groups of TSSs for initiating transcription. Consequently, it would be more accurate to refer to this occurrence as ‘*transcription start site clusters (TSSCs)*’ (49, 50). Our study on KSHV corroborates the finding that transcription initiation represents a collection of bases. We found that this phenomenon could also be extended to transcription termination (**Supplemental Figure 3**). However, for the sake of simplicity, we employ the labels ‘TSS’ and ‘TES’, referring to the aggregate of individual TSSs or TESs within a short sequence. It is still unclear whether this variation has a functional role or if it just represents transcriptional noise. Many of the detected end positions may also be of technical origin (51).

### Characterization of the viral promoters

In this part of the study, we identified the promoter elements using SeqTools, an in-house software. Specifically, we detected 81 TATA boxes, on average 31.23 bps upstream from the TSSs, and 116 GC boxes on average 69.03 bps upstream from the TSSs (**Supplemental Table 1**) using this pipeline. TSSs with a TATA box are rich in G/A bases at their first positions, while C/T base richness is observed at position −1 (**Figure 4**). TSSs without a TATA box are G-rich both at the 0 and +1 positions. We identified 20 CAAT boxes on average 117.35 bps away from the TSS. From the transcription factor IIB binding sites, we identified two sets of B recognition elements (BREs) proximately upstream of the TATA boxes: (1) 4 BREus on average 38.25 bps away from the TSS, and (2) 33 BREds on average 23.66 bps away from the TSS. We identified two downstream promoter elements (DPEs) on average 28 bps from the TSSs. In total, we identified 557 TSSs. Among these, 470 were confirmed by dRNA-Seq, 322 by CAGE sequencing, and the RAMPAGE data confirmed 190 TSSs (**Supplemental Table 1)**.

**Figure 4.**
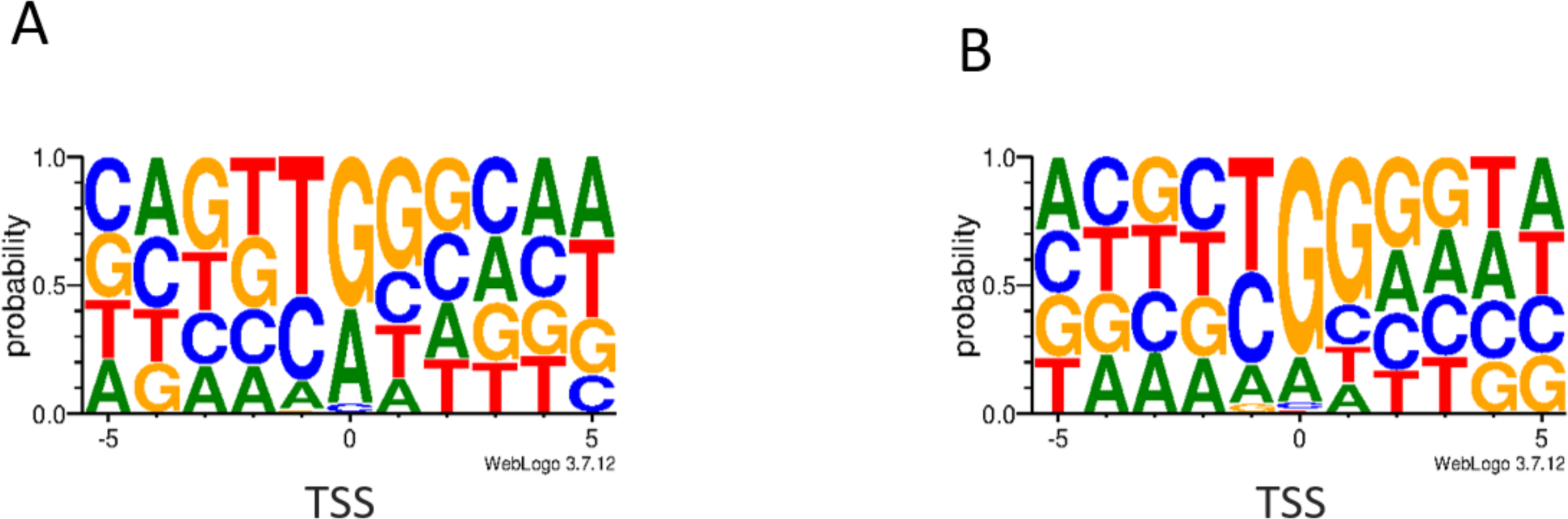
TSS consensus sequences identified in KSHV. The nucleotide composition of TSSs was identified in this study. **(A)** Genomic surrounding of TSSs with TATA box within a ± 5 bp interval. The first letter of TSSs (position 0) is enriched with G/A bases. **(B)** Genomic surrounding of TSSs without TATA box within a ± 5 bp interval. The 0 and +1 TSS positions are enriched with G letters. Base frequencies are depicted by WebLogo.

### Transcriptional activity along the KSHV genome

Unprocessed sequencing reads from both dcDNA-Seq and dRNA-Seq were used to display the transcriptional activity throughout the entire KSHV genome (**Figure 5**). This method enables the collection of a maximal number of transcription reads prior to any bioinformatic filtering. Given that LRS has a bias towards short reads, the per-point coverage is certainly not entirely accurate. Yet, it is noticeable that transcriptional activity spans the full length of the viral genome. It is assumed that with increased data coverage, transcriptional activity from every base on both DNA strands could be detected.

**Figure 5.**
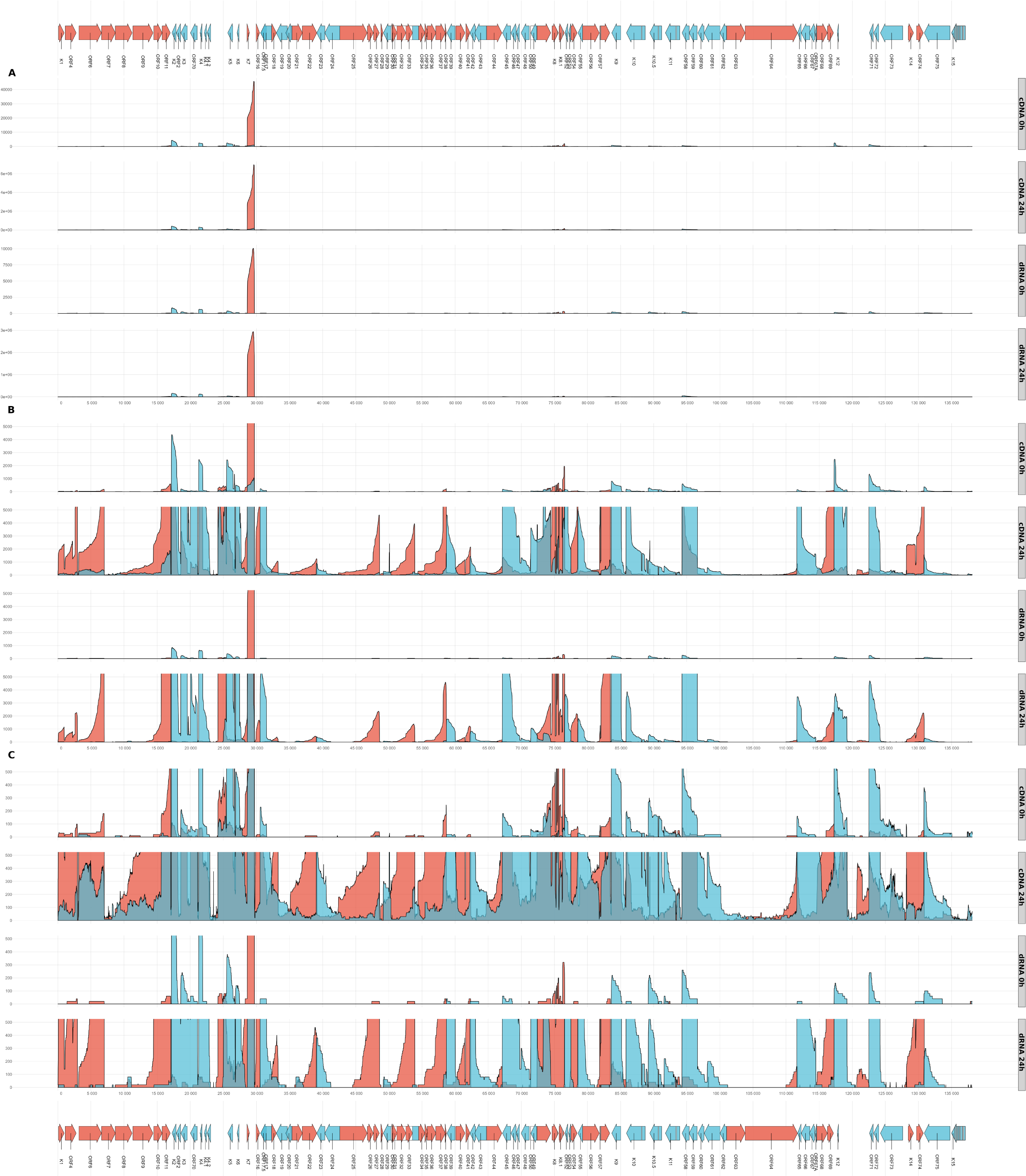
Transcriptional activity along the KSHV genome. This figure shows the dcDNA and dRNA sequencing datasets derived from the 0 hpi and 24 hpi samples. The value associated with each nucleotide was determined by tallying the number of reads overlapping that specific position. In the dcDNA-Seq, only those reads were considered where the orientation was discernible via the detection of either 5’ or 3’ adapters. The data from the three replicates of dcDNA-Seq were merged. These numbers were then grouped in 100-nt intervals to form the distributions displayed. Panel (A) restricts its y-axis to auto scale, while panel (B) sets a limit of 5 000 reads, and panel (C) sets a limit of 500 reads. Gene orientations are differentiated by color: the positive strand is represented in red, and the negative strand in blue.

### Canonical mRNAs

We performed annotation of the canonical mRNAs, which are defined as the most abundant RNA isoforms expressed by protein-coding genes (**Figure 6**; **Supplemental Table 2**). Notably, not all canonical transcripts were identified when subjected to our strictest set of conditions, which were the presence in three parallel cDNA samples, detection via dRNA-Seq, CAGE-Seq and RAMPAGE-Seq, as well as the presence of an upstream promoter. Consequently, we had to loosen some of the above constraints for the annotation of the missing transcripts. However, in some viral genes, no complete RNA molecules were detected at all, even under the less stringent criteria, probably due to the low level of gene expression (e.g. ORF7), and the large size of the transcript (e.g. ORF63, ORF64).

**Figure 6.**
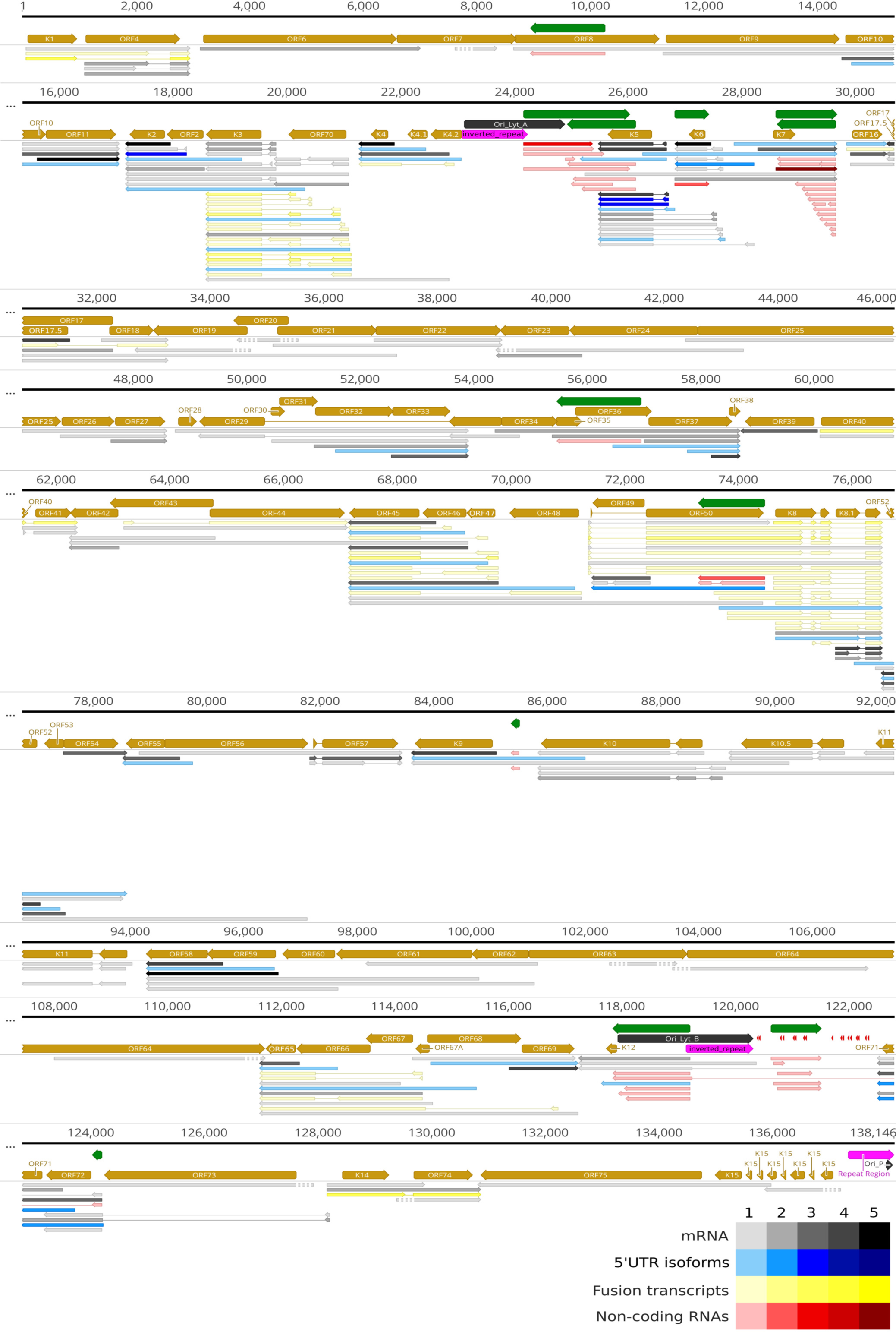
Canonical KSHV transcripts. Canonical mRNAs, defined as the most abundant RNA isoforms, are represented by black/gray arrows, while the other transcript isoforms are shown as blue arrows. Fusion transcripts are depicted by yellow arrows. The non-coding transcripts are symbolized with red arrows. The miRNAs are shown by red arrowheads. Green arrows represent asRNAs. For those transcripts where no complete RNA molecules were detected, the absent portions of the RNA molecules were denoted with striped lines. The relative abundance of viral transcripts is denoted by varying shades: 1: 1-9 reads, 2: 20-49 reads, 3: 50-199 reads, 4: 200-999 reads, 5: >1000 reads.

### Non-coding transcripts

We have also identified lncRNAs and pre-miRNAs (**Figure 6**). Non-coding transcripts possess their own promoters and are located in intragenic (such as LAN), or intergenic positions, or they overlap mRNAs in an antiparallel manner [antisense RNAs (asRNAs)]. However, antiparallel segments at either end of the mRNAs can also be found. This occurs when nearby genes on opposite strands overlap in their transcriptional activity, either convergently or divergently. Transcript isoforms that have exceptionally lengthy 5′ untranslated region (UTR) may also be ncRNAs, given the substantial gap between the TSS and translation initiation sites (TISs), Despite the ambiguity in this matter, we depict them as protein-coding transcripts in **Figure 6**. In addition to the previously published ncRNAs, we were able to identify new transcript isoforms of this type (**Supplemental Table 2**).

### Spliced transcripts

For the identification of splice sites within the KSHV transcripts, we employed highly stringent criteria: we not only required their occurrence in a minimum of three different samples, but we also mandated the presence of splice donor and acceptor consensus sites (GT/AG) and detection by dRNA-Seq in addition to direct cDNA sequencing. The dRNA-Seq validation process included checking either splice sites or the entire RNA sequences. It is known that poxvirus transcripts do not undergo splicing (52). Therefore, alongside KSHV, we also performed dcDNA-Seq and dRNA-Seq using the monkeypox transcripts to determine whether the library preparation, or the sequencing protocols might yield false splicing. Our LoRTIA software did not identify any potential introns, thus confirming that the spliced transcripts we observed in KSHV are not artifacts but rather of biological origin (supporting data not shown). We assume that the discarded, low-abundance spliced reads are also genuine spliced transcripts, even though they might not have a function. Furthermore, we noticed significant heterogeneity in the splicing sites, wherein the same splice donor site was paired with different acceptor sites, and vice versa (**Supplemental Table 1**). Another notable characteristic is that splice consensus sites are surrounded by numerous alternative splice sites in their proximity. Our annotated introns were compared to already described introns published by others (**Supplemental Figure 2A**). Here we listed 35 putative, hitherto undetected splice junctions based on dcDNA-Seq and dRNA-Seq (**Supplemental Table 1**). Fusion transcripts represent spliced sequences derived from at least two neighboring or proximate genes, encompassing chimeric UTRs or coding regions. In principle, the downstream partner in ORF fusions can be positioned in-frame or out-of-frame. In this work, we identified 10 genomic regions encoding fusion transcripts (**Figure 6**).

### Identifying 5’ and 3’ UTR isoforms of mRNAs

Determining the exact boundaries of 5′ UTRs of RNAs represents a pivotal part of transcript annotation. Methods, such as Cap-selection, Terminase enzyme utilization, CAGE-Seq, 5′RACE, and RAMPAGE-Seq analyses have limitations for validating particular TSSs with high degree of uncertainty. This issue presents a notable problem particularly for reads with low abundance and also for shorter reads (roughly 300-1,500 base pairs), which appear to be more frequent in raw data due to the aforementioned biases. Therefore, we implemented even more stringent criteria for the annotation of these reads, which were as follows: we have raised the score cutoff for CAGE-Seq data to 50 counts. The likely consequence of our strict criteria system is that numerous low-abundance transcripts remained unidentified, e.g. mRNAs expressed from ORF7, ORF19, ORF20, ORF63, ORF64, and K15 Our findings indicated that the average length of 5′ UTRs in canonical transcripts was 549.66 bp (median 257 bp, SD=746.46), whereas the mean length of 3′ UTRs of canonical transcripts was 188.83 bp (median 78 bp, SD=353.12). It should be mentioned that the potential nested mRNAs with truncated in-frame ORFs (ifORFs) are not regarded as transcript isoforms. Given that these RNAs produce distinct protein molecules, they are addressed in the subsequent section.

### Non-canonical ORFs with putative coding function

Nested genes reside within the coding regions of host genes. They share the same TES but possess different TSSs compared to canonical transcripts. Nested genes have shorter ifORFs, which if translated, would encode N-terminally truncated polypeptides. The existence of this transcript category is well-documented in herpesviruses (29), which has also been identified in KSHV (53). To find evidence for the translation of novel ifORFs, we reanalyzed the RiboSeq data published by others (41) (**Supplemental Figures 4 and 5**). The latter study (41) detected several upstream ORFs (uORF), short ORFs (sORF), and some internal ORFs (intORF). To gauge the total protein-coding capacity of certain KSHV transcripts, we first identified all potential ORFs in the full-length viral mRNAs. We found 19,930 potential ORFs on the annotated transcripts that include either cognate or non-cognate start sites (**Supplemental Figures 6 and 7A**). Only those transcripts were selected that contained an internal, co-terminal ORF (altogether 24 ifORFs). Based on our analysis, we identified 24 ifORFs that contained annotated truncated ORFs: K10, K2, K3, K4, K5, K6, K9, ORF11, ORF16, ORF17, ORF17.5, ORF22, ORF27, ORF39, ORF45, ORF49, ORF50, ORF54, ORF55, ORF57, ORF58, ORF6, ORF69 and ORF70 (**Supplemental Figure 7B**).

**Figure 7.**
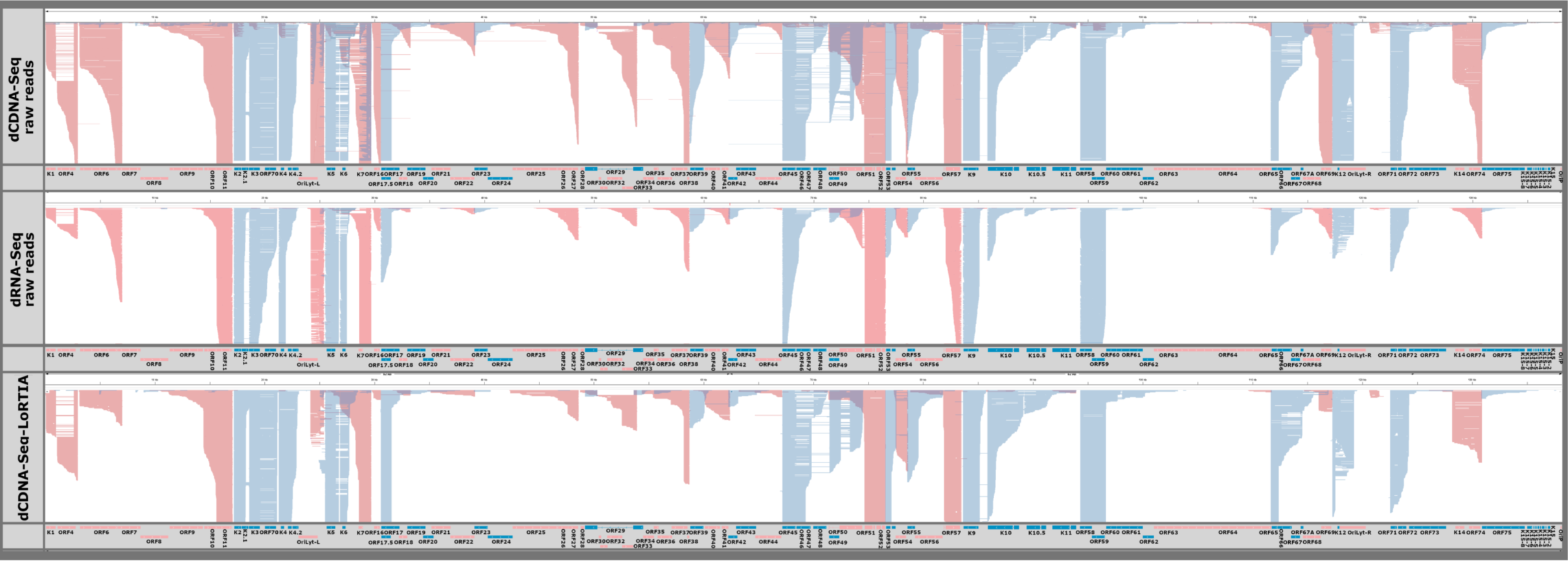
Transcriptional overlaps. The diagram displays raw reads for both dcDNA and dRNA without any added selection criteria (upper panels) alongside the genome. Meanwhile, the lower panel showcases reads that passed the quality filtering by LoRTIA and possess the correct adapters. This illustration highlights the pronounced transcriptional overlaps, notably between transcripts oriented divergently and to a lesser extent, convergently. Genes and RNAs transcribed from left to right (on the + strand) are marked in red, while those transcribed from right to left (on the − strand) are colored blue.

Subsequently, we reanalyzed the raw RiboSeq data published by others (41) aligning them with the 5’-truncated transcript data from our LRS approach to obtain the translationally active sites on the viral mRNAs. This led us to identify multiple hidden translation initiation sites (TISs) within the canonical ORFs. We chose only the highest significant TISs (p<0.05) and associated them with the potential ifORFs of K2, K6, ORF11, ORF17, ORF17.5, ORF45, and ORF57. These genes use predominantly the canonical AUG and to a lesser extent the AUA non-canonical start codons and are terminated in-frame compared to the given ORF. Our algorithm also detected TIS-peaks on the transcript isoforms of K3, K4, K5, K9, ORF16, ORF22, ORF27, ORF39, ORF49, ORF50, ORF54, ORF55, ORF58, ORF6, ORF69 and ORF70 with notably high occurrence (however, their z-score significance values were slightly above the p-value at 24 hpi (**Supplemental Table 2. Supplemental Figure 7C**).

Six intORFs have previously been identified in KSHV (41). Yet, based on our LRS data, three of them would be more aptly termed as uORFs due to their short length and the presence of a longer downstream ORF within the same transcript. We determined that the intORF within ORF10 is not transcribed separately but rather in conjunction with the downstream ORF11, resulting in a 5’UTR variant. Similarly, the intORF of K8.1 seems to act as an uORF for the transcript associated with the second exon of the K8.1 gene, which is transcribed independently (**Supplemental File 1).**

### Coding potential of ‘non-coding’ KSHV sequences

Another question we also explored is whether the long ncRNAs of KSHV possess any coding potential. We detected asRNAs in the genomic region encoding IE genes and other tandem areas of the KSHV genome. Previously, these regions were believed to be non-coding. To assess the coding capability of these genomic segments, we re-examined the original RiboSeq data (41) and identified the ribosomal footprints on the KSHV transcripts. Additionally, a proteomic study of lytic KSHV infection employing RNA-tiling array and protein LS-MS/MS method had been conducted (54). By linking these datasets, we were able to associate with viral peptide sequences that had no previously identified corresponding mRNAs. In-depth reanalysis of these proteomic datasets revealed that a portion of these unlinked viral peptides may be encoded by the asRNAs and complex transcripts that are multigenic RNAs containing at least one ORF in an opposite orientation (**Supplemental Table 3**).

### An OriLyt-spanning protein coding transcript

We identified a cluster of replication-associated RNAs (raRNAs) (55) overlapping the KSHV OriLyt-L with a shared TSS at the genomic location 24,178. This position is adjacent to a TSS previously documented by others (41, 56). Moreover, this locus contains a TATA-like element 29 bp upstream of the TSS (57). They have been categorized as non-coding in our first survey. However, in all six independently analyzed RiboSeq samples (41), a pronounced TIS peak is observed at the intergenic repeat region, positioned at the genomic location 24,214, which lies between K5 and K4.2, suggesting the existence of a novel, OriL-spanning sequence with the potential for encoding a 233 amino-acid long polypeptide. Furthermore, four peptide sequences from the KSHV proteomic dataset (54) are mapped to this intergenic area (**Supplemental Table 3**). These peptides are aligned in-frame with the recognized sORF that are terminated at genomic position 24,915 (**Supplemental Figure 8**). A search in the database of conserved protein domains, utilizing the *in silico* translated sequence of the 233 amino acids, showed a subtle similarity to eukaryotic PolyA-binding protein superfamily (3.97 e-4) and to DNA Polymerase III subunits gamma and tau (2.43 e-8). A BLASTP query revealed the highest resemblance to either hypothetical or not yet characterized proteins (with bit scores of 132 and 102) and mucin-like proteins (scoring 112).

### Transcriptional overlaps

Gene pairs can adopt parallel (→→), convergent (→←), or divergent (←→) orientations. If the canonical transcripts of gene pairs form transcriptional overlaps (TOs) with each other, we refer to it as a ’hard’ overlap. On the other hand, a ’soft’ TO occurs when only the longer TSS or TES variants form overlaps, but not the canonical transcripts. **Figure 3C** (along with **Figure 7**) and **Supplemental Table 4** shows that out of 13 canonical transcripts encoded by convergent gene pairs, 8 form hard TOs, while the remaining transcripts produce varying degrees of soft TOs. Divergent gene pairs predominantly generate hard head-to-head TOs (10 out of 12 gene pairs), with 2 instances of soft TOs. Notably, both convergent and divergent gene pairs with hard TOs exhibit more extensive overlaps due to transcriptional read-through or the generation of long TSS isoforms, respectively. Furthermore, we observed that in co-oriented viral genes, the TSS of downstream genes is in many cases located within the ORF of the upstream gene. It is important to mention that our data underrepresent divergent TOs because a significant proportion of TSSs are not detected in transcription reads due to sequencing biases favoring short sequences. Similarly, the extent of convergent overlaps is also underestimated because RNA polymerase tends to read through the poly(A) site, resulting in an extended nucleic acid stretch that gets later truncated at the TES.

### Viral transcripts expressed during latency

We have also analyzed the viral gene expression in non-induced (latent) iBCBL1-3xFLAG-RTA cells. While we were able to identify the four latency transcripts, we also detected viral transcripts that are typically expressed during the lytic phase, such as the PAN RNA whose amount was significantly higher compared to other viral RNAs (**Figure 5**). This observation is attributed to the spontaneous reactivation of the virus in a number of latently infected cells. Indeed, Landis and colleagues employing single-cell RNA sequencing to study KSHV latency detected such variability in gene expression within the latently infected PEL cell population (58).

## DISCUSSION

Over the last couple of years, sequencing technologies have significantly advanced. This rapid development, spearheaded by third-generation LRS techniques, instigated a fundamental transformation in the study of cellular and viral transcriptomes (29, 59–63). Investigations have demonstrated that the transcriptomic composition of viruses is considerably more intricate than initially thought (64). A broad array of overlapping transcripts has been unveiled in every herpesvirus family (55, 65).

Through the creation of a LRS dataset and the application of a robust transcript annotation pipeline, we effectively annotated numerous novel TSSs, TESs, introns and transcripts, and confirmed or amended previously annotated KSHV RNAs in the lytic phase (24 hpi) and during latency. We successfully identified long 5′UTR isoforms, complex transcripts, various alternative splice variants, and chimeric RNA in the KSHV transcriptome. Our splice detection results from dRNA-Seq align with other published datasets to differing extents, with only a restricted subset of introns (19) being common across all the examined data (**Supplemental Figure 2A**).

Our research significantly expanded the number of KSHV TSS and TES in comparison to previously published data (**Supplemental Figures 2B and C, Supplemental File 1**). Although LRS techniques have made advancements in accuracy, the reliable end-to-end identification of transcripts continues to pose a challenge (51). To guarantee the credibility of our results, we applied strict filtering standards and cross-checked our discoveries with validated data from additional research studies. We employed the LoRTIA pipeline, a tool developed in our lab, for the annotation of KSHV transcripts. To provide further validation of our findings, we utilized CAGE analysis, along with RAMPAGE, 3’RACE and RiboSeq data generated by other researchers. It is important to note that our strict filtering may have resulted in the loss of several rare *bona fide* transcripts.

Our research also uncovered an incredibly intricate network of TOs. Herpesvirus gene clusters comprising co-oriented genes are known to produce parallel transcriptional overlaps due to their shared transcription termination. Our approach revealed that most convergent and divergent gene pairs create ’hard’ overlaps, where their canonical transcripts overlap with each other.

Furthermore, our findings indicate that the entire KSHV genome is transcriptionally active, including both DNA strands as also shown by previous studies. Nevertheless, the question arises whether all these transcripts serve functional roles, or whether some of them are the results of a transcriptional interference mechanism (64), or perhaps, they are just transcriptional noise (66).

We identified several new nested transcripts that include co-terminal intORFs. We define ‘ifORF’ as a form of the intORF, which is detected within monocistronic transcripts in in-frame position. For ifORFs located within the 5′UTR of RNAs encoded by the upstream genes in a multicistronic transcript, it is difficult to determine whether they are just long 5′UTR isoforms of the downstream gene, or if these ifORFs are actually translated. It is an important issue, since in the latter scenario, the downstream gene would not undergo translation due to the absence of requested sequences and molecular mechanisms that would allow this process. Whether these putative 5′ truncated ORFs have true coding capacity and are all biological products remain to be determined.

The initiation codons for ifORFs are substantiated by notable TIS signals. ORF57 is one of the essential regulatory KSHV genes, which encodes the IE63 homologue of HSV-1 (ICP27). We confirmed the long isoform of its transcripts described originally by Northern-blot analysis (25) and found several internal, short isoforms as well. The K3 and K5 gene products are involved in the modulation of host cell’s immune response (67). The K3 gene possesses an intORF (41), and we successfully associated it with multiple transcript isoforms. Despite the many nested transcripts identified within K5, we did not detect ‘strong’ TIS signals, which indicates that not all embedded transcripts function as potential protein-coding mRNA.

A limited number of factors orchestrate the first steps of the KSHV lytic infection (68). We detected the highest TSS variability of transcript isoforms in ORF50-(ORF49)-K8-K8.1 locus, and the highest intron variability in the K2-ORF2-K3-ORF70 genomic region (**Supplemental File 1, Supplemental Figure 4**) involved in immune evasion.

The OriLyt-L region of KSHV holds significant interest due to the existence of repeats that provide a binding site for RTA. Transcriptional activity and TES have already been detected in this region by several independent experiments (41, 48, 54, 56, 69). However, they could not determine the exact structure of these mRNAs. In this region, the K4.1 mRNA and another labeled as T1.5 was identified using Northern-blot (56, 69). Close to the 3’end of these transcripts, a short potential protein, termed OLAP, has been previously predicted (57) and a TIS signal was detected (41). Here, we not only identified these transcripts but also discovered a subset that includes a new, extended isoform of an OriLyt-L-spanning RNA. The ORF identified here is also corroborated by RiboSeq and LS-MS/MS data (41, 54). However, we were unable to identify the 6-kb long, OriLyt-L spanning transcript T6.1 (57).

Several KSHV asRNAs have previously been identified (54, 57, 70). Notably, the 10-kbp long antisense latency transcript (ALT) might play a role as a primary regulator of OriLyt-R latency locus. Contrary to the earlier hypothesis suggesting that ALT exists without other isoforms (71), we identified shorter versions of ALT. Moreover, we also discovered other asRNA isoforms to which we could associate viral peptide fragments that were previously not assigned to any viral transcripts (54). This led us to propose that these asRNAs might be the original source for these peptides. Although peptides encoded by antisense RNAs have been observed in KSHV (72) and other herpesviruses (73, 74), their precise function continues to be an area of active debate (75).

Gammaherpesviral ncRNAs modulate host immune reactions through various mechanisms (76, 77). Among ncRNAs, the newly characterized circRNAs are produced by back-splicing (18, 78). The new introns identified in our study might serve as a source for circRNAs. The full coding capacity of the KSHV transcriptome still requires deeper exploration. While the LC-MS/MS method has challenges in detecting low-abundance proteins, employing advanced protein sequencing methods on nanopore arrays could address this issue (79, 80).

Taken together, the information generated by integrating the data obtained from our combined methods has expanded our understanding of the viral transcriptome architecture of KSHV. Our results illustrate that the lytic transcriptome of the KSHV is even more complex than it was initially described. Our study underscores the significance of utilizing a combined multiplatform strategy in transcriptomics.

## MATERIALS AND METHODS

### Cell culture, RT-qPCR and immunoblot analyses

The iBCBL1-3xFLAG-RTA cell line, a KSHV-positive primary effusion lymphoma (PEL) (81), was grown in RPMI 1640 medium supplemented with 10% Tet System Approved FBS (TaKaRa), penicillin/streptomycin, and 20 μg/ml of hygromycin B. To initiate KSHV lytic reactivation, one million iBCBL1-3xFLAG-RTA cells were treated with 1 μg/ml of doxycycline (Dox) for a 24-hour period. For assessing KSHV gene expression via RT-qPCR, cDNA was produced using the iScript cDNA Synthesis kit (Bio-Rad), followed by SYBR green-based real-time quantitative PCR analysis with gene-specific primers. The relative expression of viral genes was determined using the delta-delta Ct method, with 18S serving as the normalization factor. The primer sequences were previously reported by Toth et al. (82). The immunoblots employed the following antibodies: anti-FLAG (Sigma F1804), anti-ORF6 (provided by Dr. Gary S. Hayward from Johns Hopkins University), and anti-Tubulin (Sigma T5326).

### RNA isolation, poly(A) selection, and measurement of nucleic acid quality and quantity

**Supplemental Text** contains the comprehensive protocols.

### Direct cDNA sequencing

Libraries were generated from the poly(A)^+^ RNA fractions without PCR amplification using the ONT’s Direct cDNA Sequencing Kit (SQK-DCS109) following the ONT manual. In summary, RNAs were mixed with ONT VN primer and 10 mM dNTPs and incubated at 65°C for 5 minutes. Next, RNaseOUT (Thermo Fisher Scientific), 5x RT Buffer (Thermo Fisher Scientific), and ONT Strand-Switching Primer were added to the mixtures, which were then incubated at 42°C for 2 minutes. The Maxima H Minus Reverse Transcriptase enzyme (Thermo Fisher Scientific) was added to the samples to create the first cDNA strands. The reaction took place at 42°C for 90 minutes, and the reactions were stopped by heating the samples at 85°C for 5 minutes. The RNAs from the RNA: cDNA hybrids were eliminated using the RNase Cocktail Enzyme Mix (Thermo Fisher Scientific) in a 10-minute reaction at 37°C.

The second strand of cDNAs was synthesized using LongAmp Taq Master Mix [New England Biolabs (NEB)] and ONT PR2 Primer. The PCR condition applied was: 1 minute at 94°C, 1 minute at 50°C, and 15 minutes at 65°C. Subsequently, end-repair and dA-tailing were performed with the NEBNext End repair/dA-tailing Module (NEB) reagents at 20°C for 5 minutes, followed by heating the samples at 65°C for 5 minutes. Adapter ligation was conducted using the NEB Blunt/TA Ligase Master Mix (NEB) at room temperature for 10 minutes. The ONT Native Barcoding (12) Kit was employed to label the libraries, which were then loaded onto ONT R9.4.1 SpotON Flow Cells (200 fmol mixture of libraries was loaded onto one flow cell). AMPure XP Beads were utilized after each enzymatic step, and samples were eluted in UltraPure™ nuclease-free water (Invitrogen).

### Native RNA sequencing

For the dRNA-seq experiments, an RNA mixture (pooled biological replicates) was prepared, which included RNA from Poly(A)+ samples. The T10 adapter containing oligo dT for RT priming and the RNA CS for monitoring sequencing quality (both from the ONT kit) were added to the RNA mix, along with NEBNext Quick Ligation Reaction Buffer and T4 DNA ligase (both from NEB). The reaction incubated for 10 minutes at room temperature. Subsequently, 5x first-strand buffer, DTT (both from Invitrogen), dNTPs (NEB), and UltraPure™ DNase/RNase-Free water (Invitrogen) were added to the samples. Finally, the SuperScript III enzyme (Thermo Fisher Scientific) was combined with the sample, and the RT reaction was conducted at 50°C for 50 minutes, followed by enzyme heat inactivation at 70°C for 10 minutes.

The RNA adapter (from the ONT kit) was ligated to the RNA: cDNA hybrid sample using the NEBNext Quick Ligation Reaction Buffer and T4 DNA ligase at room temperature for 10 minutes. RNAClean XP Beads were employed after each subsequent enzymatic step. Two flow cells were utilized for dRNA-seq, with 100 fmol of the sample loaded onto each.

### Cap Analysis of Gene Expression (CAGE)

The CAGE™ Preparation Kit (DNAFORM, Japan) was employed according to the manufacturer’s guidelines (see **Supplemental Text** for the details).

### ONT sequencing data analysis

The Guppy software (v3.4.5) was utilized for basecalling ONT-MinION sequencing reads. Reads with a quality filter of 8 (default) were mapped to the reference genome using minimap2, applying settings: -ax splice -Y -C5 -cs. Mapping statistics were calculated using the ReadStatistics script from Seqtools (https://github.com/moldovannorbert/seqtools). The LoRTIA toolkit (alpha version, accessed on 20 August 2019, https://github.com/zsolt-balazs/LoRTIA) was employed for identifying TESs, TSSs, and introns, and reconstructing transcripts based on these features. The LoRTIA workflow with default settings included: 1) for dRNA and dcDNA sequencing: −5 TGCCATTAGGCCGGG --five_score 16 -- check_in_soft 15 −3 AAAAAAAAAAAAAAA --three_score 16 s Poisson–f true; and 2) for o(dT)-primed cDNA reads: −5 GCTGATATTGCTGGG --five_score 16 --check_in_soft 15 −3 AAAAAAAAAAAAAAA --three_score 16 s Poisson–f true.

A read was accepted if the adapters were accurate, polyA tails were present and no false priming events were detected by LoRTIA. For introns, only those present in dRNA sequencing were accepted, as this method is regarded as the ‘Gold Standard’ for identifying alternative splicing events. Some transcripts were manually included if they were a long TSS variant of already accepted TSSs. MotifFinder from Seqtools was employed to find promoter elements around the accepted TSSs.

### Illumina CAGE sequencing data analysis

Read quality was assessed using FastQC (https://www.bioinformatics.babraham.ac.uk/projects/fastqc). Reads were trimmed using TrimGalore (https://github.com/FelixKrueger/TrimGalore) with the following settings: -length 151 -q. The STAR aligner (version 2.7.3.a) was employed to map the reads to the KSHV strain TREx reference genome (GQ994935.1), utilizing --genomeSAindexNbases 8 and default parameters. The CAGEfightR R package was used to identify TSSs and TSS clusters with a minimum pooled value cutoff of 0.1 (pooledcutoff=0.1).

### Downstream data analysis and visualization

Data analysis downstream was conducted and figures were created as described in **Supplemental text**.

### Data availability

Basecalled sequencing FastQ files used in this study have been deposited to in European Nucleotide Archive (ENA) under the following Bioproject accession number: PRJEB60022 and link: https://www.ebi.ac.uk/ena/browser/view/PRJEB60022. Accession numbers and statistics of files containing the CAGE and MinION mapped reads are summarized in **Supplemental Table 5**.

### Code availability

The LoRTIA software suite and the SeqTools are available on GitHub.

LoRTIA: https://github.com/zsolt-balazs/LoRTIA

R scripts: https://github.com/Balays/Rlyeh

R workflow: https://github.com/Balays/KSHV_RNASeq

## Acknowledgments

This research was supported by National Research, Development and Innovation Office (NRDIO), Researcher-initiated research projects (Grant numbers: K 128247 and K 142674) to ZB and by the NRDIO Research projects initiated by young researchers (Grant number: FK 142674) to DT. The work was also supported by NIH grant R01AI132554 to ZT. IP was supported by UNKP-22-4 - SZTE-310. The APC fee was covered by the University of Szeged, Open Access Fund: 6049. We are thankful to Carolina Arias for sharing the RiboSeq raw data.

## Ethics declarations

Not applicable

## Author contributions

**ÁD**: Participated in RNA purification and long-read sequencing

**ÁF**: Contributed to bioinformatics and data interpretation

**BK:** Contributed to bioinformatics, data interpretation and visualization

**DT**: Performed library preparation, long-read and CAGE sequencing, participated in data interpretation, and drafted the manuscript

**GG**: Contributed to sequencing and bioinformatics

**GT**: Conducted bioinformatics, data, interpretation, and integration of data

**IP**: Participated in data analysis integration, visualization and drafted the manuscript

**LMS**: Participated in cell propagation and carried out RNA purification

**ZB**: Conceived and designed the experiments, supervised the study, and wrote the manuscript

**ZT**: Contributed to the experiment design and drafted the manuscript All authors read and approved the final paper.

## SUPPLEMENTAL MATERIAL

### Supplemental Figures

**Supplemental Figure 1. Transcriptional end sites of KSHV RNAs**

In this figure, the TES distribution aligned with the KSHV genome annotation is illustrated, highlighting each ORF. Panels A and B illustrate the coverage from the dcDNA and dRNA sequencing datasets derived from the 24 hpi samples, respectively. For each nucleotide the TES signal strength value was calculated by counting the reads that have their 3’ ends aligning with that specific position. In the context of the dcDNA-Seq, reads were only included if their orientation could be identified through the presence of either 5’ or 3’ adapters. The dcDNA-Seq data of the three replicates were merged. Subsequently, the TES signal strength values were clustered in 100-nt intervals to represent the distributions. The y-axis in panel (A) tops out at 2000 reads, whereas in panel (B) it is limited to 200 reads. Gene orientations are color-coded: red for the positive strand and blue for the negative strand.

**Supplemental Figure 2. Introns, TSSs and TESs of KSHV transcripts**

**A.** The Venn-diagram displays the count of introns sourced from published data and compares them with our findings obtained through dRNA and dcDNA sequencing.

**B.** The onion-diagram illustrates the cumulative count of annotated 5′ read ends utilized in the transcript assembly as TSS. End positions of reads from dRNA and CAGE samples were counted within a +/-10 bp window.

**C.** The oligo(dT)-based dcDNA-Seq and dRNA-Seq yield comparable 3′ ends for mRNAs.

**Supplemental Figure 3. Heterogeneity of transcription initiation and termination**

The figure is composed of six panels, illustrating a wide variety of TSS (A1, B1 and C1) and TES (A2, B2 and C2) distributions along three genomic regions. The upper panel (A1-A2) shows the K2-ORF2 genomic region, the middle panel shows the region around K4 (B1-B2), while the bottom panel (C1-C2) shows the K7-PAN region. The x-axes show the genomic position, while the y-axes represent the number of either 5-prime ends, or 3-prime ends of reads in that position for the TSS or TES panels, respectively. For each panel, there are three sub-panels in different scales: the leftmost panels have no y-axis limit, the middle panels have a limit of 500 reads, while the rightmost panels have 50 reads. Genes depicted on the figure show examples for both TSS and TES clustering, which phenomenon is independent from the sequencing method.

**Supplemental Figure 4. Total transcriptome of KHSV**

Canonical mRNAs, defined as the most abundant RNA isoforms, are represented by black/gray arrows, while the other transcript isoforms are shown as blue arrows. The non-coding transcripts are symbolized with red arrows. Truncated transcripts are indicated by orange arrows. The miRNAs are shown by red arrowheads. Green arrows represent asRNAs. For RNA molecules without detected complete sequences, the missing sections of the transcripts were indicated with striped lines. The relative abundance of viral transcripts is denoted by varying shades: 1: 1-9 reads, 2: 20-49 reads, 3: 50-199 reads, 4: 200-999 reads, 5: >1000 reads.

**Supplemental Figure 5. Comparison of TIS and TSS distribution along the entire viral genome**

Panels (A) and (B) display the TSS distribution from CAGE-Seq and the TIS distribution from RiboSeq, respectively. For each nucleotide a signal strength value was calculated by counting the reads with their 5’ ends at that position. These values were subsequently summed within 100-nt windows to produce the displayed distributions. For panel (A), the y-axis is capped at 1000 counts for CAGE-Seq and 5000 counts for RiboSeq. In panel (B), the limits are 1000 counts for CAGE-Seq and 5000 counts for RiboSeq. The distributions are the KSHV genome annotations with ORFs shown beneath. Gene orientations are color-coded: red for the positive strand and blue for the negative strand.

**Supplemental Figure 6. The coding capacity of KSHV**

The figure shows the widths of all the predicted ORFs in the KSHV genome according to their start codons. Each point represents a single ORF, with the y-axis showing its width (nt) and the x-axis its start codon. Points are colored with blue in the case of canonical ORFs (all of them starting with ATG), otherwise they are colored with light brown. The upper panel (A) shows those ORFs that are coterminal with a canonical ORF, while the lower panel (B) shows those that are not.

**Supplemental Figure 7. Ribosome footprint signals on 5′-truncated transcripts**

Panel **(A)** shows the potential coding capacity of the different annotated transcripts according to their types, including mono-, bi-, polycistronic, and 5′ truncated transcripts.

Significant RiboSeq footprint signals on 5′-truncated transcripts in the 24h CHX RiboSeq sample. Panel **(B)** shows the signal strength of the predicted ifORFs encoded by the truncated transcripts. Each point represents a single predicted ORF for each transcript. The x-axis shows the name of the host ORF, while the y-axis shows the ratio of the TIS strength around the predicted ORF, and that of its host ORF. The TIS signal for each in-frame ORF was calculated as the sum of the read counts around +/− 2 nts of its start position. Panel (**C)** shows the result of the significant TIS peak detection on the 5′ truncated transcripts. Each point represents a TIS peak. The x-axis shows the canonical ORFs, while the z-score of each TIS peaks are depicted on the y-axis. Red points represent peaks with insignificant, while blue points represent peaks with significant p-values (significance cutoff = 0.05).

**Supplemental figure 8. A novel OriLyt-spanning protein-coding transcript**

We found that a group of transcripts overlaps the direct repeats at the intergenic region of OriLyt-L (indicated by green color). Annotated features, like promoter, TSS and TES are marked as well as footprints of the Ribo-Seq signals obtained by Arias and colleagues (41) (upper panel). Peptides from data published by Dresang colleagues. (54) are marked as green arrows.

### Supplemental Tables

**Supplemental Table 1. Comparing our KSHV TSS and TES data with findings from other publications**

A: TSS: Here, we list all TSS positions (sourced from our cDNA samples) that met our criteria based on the TSS annotation (detailed in the Methods section). Their alignment with CAGE, RAMPAGE, and dRNA coordinates is indicated. Additionally, any identified cis-regulatory elements are also included in this table.

B: TES: In this table, we list all positions that satisfied our criteria for TES annotation. The TESs presented here originate from the cDNA and dRNA samples, and any alignment with other coordinates is highlighted. Additionally, other identified polyA signals are included in this table.

C: Intron: Here, we enumerate all splice junctions according to their strand and coordinates in reference to the genome: GQ994935.1.

**Supplemental Table 2. List of the KSHV transcripts**

This table presents the genome coordinates, aligned to the reference genome GQ994935.1. The abundance of the different transcript types are as follows: A: mRNAs, B: non-coding RNAs, C: truncated transcripts, D: spliced transcripts, E: complex transcripts. For inclusion in the table, a transcript needed to appear in a minimum of two dcDNA samples, dRNA, or CAGE samples.

**Supplemental Table 3. Peptides and transcripts**

Dresang and co-workers (54) compiled the peptide sequences from their LC-MS/MS experiments that were detected in both lytic and latent infections of the BCBL-1 cell line. The table presents peptides that were associated with annotated transcripts.

**Supplemental Table 4. Transcriptional overlaps**

This table summarizes the transcriptional read-throughs between neighboring gene pairs according to their orientation: A: convergent, B: divergent, or C: parallel.

**Supplemental Table 5. Accession numbers of sequencing files**

Sequenced reads are uploaded into ENA. This table summarizes the sample accession numbers, read accession numbers and basic statistics of mapped reads: A: MinION sequencing; B: Illumina sequencing.

**Supplemental File 1. Transcriptomic landscape of KSHV.** The file contains all of the detected transcripts by this analysis complemented with previously published features of KSHV transcriptome in GFF, GenBank and in Geneious file format. Files are available from the Figshare link: https://figshare.com/account/articles/24139197

## References

1. Goncalves PH, Ziegelbauer J, Uldrick TS, Yarchoan R. 2017. Kaposi sarcoma herpesvirus-associated cancers and related diseases. Curr Opin HIV AIDS 12:47–56.

2. Chang Y, Cesarman E, Pessin MS, Lee F, Culpepper J, Knowles DM, Moore PS. 1994. Identification of Herpesvirus-Like DNA Sequences in AIDS-Associated Kaposi’s Sarcoma. Science (80) 266:1865–1869.

3. Ganem D. 2010. KSHV and the pathogenesis of Kaposi sarcoma: listening to human biology and medicine. J Clin Invest 120:939–949.

4. Cesarman E, Knowles DM. 1999. The role of Kaposi’s sarcoma-associated herpesvirus (KSHV/HHV-8) in lymphoproliferative diseases. Semin Cancer Biol 9:165–174.

5. Chandriani S, Xu Y, Ganem D. 2010. The Lytic Transcriptome of Kaposi’s Sarcoma-Associated Herpesvirus Reveals Extensive Transcription of Noncoding Regions, Including Regions Antisense to Important Genes. J Virol 84:7934–7942.

6. Uldrick TS, Whitby D. 2011. Update on KSHV epidemiology, Kaposi Sarcoma pathogenesis, and treatment of Kaposi Sarcoma. Cancer Lett 305:150–162.

7. Bechtel JT, Liang Y, Hvidding J, Ganem D. 2003. Host Range of Kaposi’s Sarcoma-Associated Herpesvirus in Cultured Cells. J Virol 77:6474–6481.

8. Speck SH, Ganem D. 2010. Viral latency and its regulation: Lessons from the γ-Herpesviruses. Cell Host Microbe. Cell Press 10.1016/j.chom.2010.06.014.

9. Kedes DH, Lagunoff M, Renne R, Ganem D. 1997. Identification of the gene encoding the major latency-associated nuclear antigen of the Kaposi’s sarcoma-associated herpesvirus. J Clin Invest 100:2606–2610.

10. Dittmer D, Lagunoff M, Renne R, Staskus K, Haase A, Ganem D. 1998. A Cluster of Latently Expressed Genes in Kaposi’s Sarcoma-Associated Herpesvirus. J Virol 72:8309–8315.

11. Cai X, Lu S, Zhang Z, Gonzalez CM, Damania B, Cullen BR. 2005. Kaposi’s sarcoma-associated herpesvirus expresses an array of viral microRNAs in latently infected cells. Proc Natl Acad Sci 102:5570–5575.

12. Samols MA, Hu J, Skalsky RL, Renne R. 2005. Cloning and Identification of a MicroRNA Cluster within the Latency-Associated Region of Kaposi’s Sarcoma-Associated Herpesvirus. J Virol 79:9301–9305.

13. Speck SH, Ganem D. 2010. Viral Latency and Its Regulation: Lessons from the γ-Herpesviruses. Cell Host Microbe 8:100–115.

14. Uppal T, Banerjee S, Sun Z, Verma S, Robertson E. 2014. KSHV LANA—The Master Regulator of KSHV Latency. Viruses 6:4961–4998.

15. Ballestas ME, Chatis PA, Kaye KM. 1999. Efficient Persistence of Extrachromosomal KSHV DNA Mediated by Latency-Associated Nuclear Antigen. Science (80-) 284:641–644.

16. Lin Y-T, Kincaid RP, Arasappan D, Dowd SE, Hunicke-Smith SP, Sullivan CS. 2010. Small RNA profiling reveals antisense transcription throughout the KSHV genome and novel small RNAs. RNA 16:1540–1558.

17. Qin J, Li W, Gao S-J, Lu C. 2017. KSHV microRNAs: Tricks of the Devil. Trends Microbiol 25:648–661.

18. Tagawa T, Gao S, Koparde VN, Gonzalez M, Spouge JL, Serquiña AP, Lurain K, Ramaswami R, Uldrick TS, Yarchoan R, Ziegelbauer JM. 2018. Discovery of Kaposi’s sarcoma herpesvirus-encoded circular RNAs and a human antiviral circular RNA. Proc Natl Acad Sci 115:12805– 12810.

19. Ungerleider NA, Jain V, Wang Y, Maness NJ, Blair R V., Alvarez X, Midkiff C, Kolson D, Bai S, Roberts C, Moss WN, Wang X, Serfecz J, Seddon M, Lehman T, Ma T, Dong Y, Renne R, Tibbetts SA, Flemington EK. 2019. Comparative Analysis of Gammaherpesvirus Circular RNA Repertoires: Conserved and Unique Viral Circular RNAs. J Virol 93.

20. Sun R, Lin S-F, Gradoville L, Yuan Y, Zhu F, Miller G. 1998. A viral gene that activates lytic cycle expression of Kaposi’s sarcoma-associated herpesvirus. Proc Natl Acad Sci 95:10866– 10871.

21. Guito J, Lukac DM. 2012. KSHV Rta Promoter Specification and Viral Reactivation. Front Microbiol 3.

22. Di C, Zheng G, Zhang Y, Tong E, Ren Y, Hong Y, Song Y, Chen R, Tan X, Yang L. 2022. RTA and LANA Competitively Regulate let-7a/RBPJ Signal to Control KSHV Replication. Front Microbiol 12.

23. Conrad NK. 2016. New insights into the expression and functions of the Kaposi’s sarcoma-associated herpesvirus long noncoding PAN RNA. Virus Res 212:53–63.

24. Sun R, Lin SF, Gradoville L, Miller G. 1996. Polyadenylated nuclear RNA encoded by Kaposi sarcoma-associated herpesvirus. Proc Natl Acad Sci 93:11883–11888.

25. Bello LJ, Davison AJ, Glenn MA, Whitehouse A, Rethmeier N, Schulz TF, Barklie Clements J. 1999. The human herpesvirus-8 ORF 57 gene and its properties. J Gen Virol 80:3207–3215.

26. Dittmer DP. 2003. Transcription profile of Kaposi’s sarcoma-associated herpesvirus in primary Kaposi’s sarcoma lesions as determined by real-time PCR arrays. Cancer Res 63:2010–5.

27. Jenner RG, Albà MM, Boshoff C, Kellam P. 2001. Kaposi’s Sarcoma-Associated Herpesvirus Latent and Lytic Gene Expression as Revealed by DNA Arrays. J Virol 75:891–902.

28. Nakamura H, Lu M, Gwack Y, Souvlis J, Zeichner SL, Jung JU. 2003. Global Changes in Kaposi’s Sarcoma-Associated Virus Gene Expression Patterns following Expression of a Tetracycline-Inducible Rta Transactivator. J Virol 77:4205–4220.

29. Tombácz D, Csabai Z, Oláh P, Balázs Z, Likó I, Zsigmond L, Sharon D, Snyder M, Boldogkői Z. 2016. Full-length isoform sequencing reveals novel transcripts and substantial transcriptional overlaps in a herpesvirus. PLoS One 11:1–29.

30. O’Grady T, Wang X, Höner zu Bentrup K, Baddoo M, Concha M, Flemington EK, Höner zu Bentrup K, Baddoo M, Concha M, Flemington EK. 2016. Global transcript structure resolution of high gene density genomes through multi-platform data integration. Nucleic Acids Res 44:1–17.

31. Balázs Z, Tombácz D, Szűcs A, Snyder M, Boldogkői Z. 2017. Long-read sequencing of the human cytomegalovirus transcriptome with the Pacific Biosciences RSII platform. Sci Data 4:170194.

32. Boldogkői Z, Szűcs A, Balázs Z, Sharon D, Snyder M, Tombácz D. 2018. Transcriptomic study of herpes simplex virus type-1 using full-length sequencing techniques. Sci Data 10.1038/sdata.2018.266.

33. Boldogkői Z, Moldován N, Szűcs A, Tombácz D. 2018. Transcriptome-wide analysis of a baculovirus using nanopore sequencing. Sci Data 5:180276.

34. Depledge DP, Srinivas KP, Sadaoka T, Bready D, Mori Y, Placantonakis DG, Mohr I, Wilson AC. 2019. Direct RNA sequencing on nanopore arrays redefines the transcriptional complexity of a viral pathogen. Nat Commun 10.

35. Cackett G, Matelska D, Sýkora M, Portugal R, Malecki M, Bähler J, Dixon L, Werner F. 2020. The African Swine Fever Virus Transcriptome. J Virol 94.

36. Tombácz D, Dörmő Á, Gulyás G, Csabai Z, Prazsák I, Kakuk B, Harangozó Á, Jankovics I, Dénes B, Boldogkői Z. 2022. High temporal resolution Nanopore sequencing dataset of SARS-CoV-2 and host cell RNAs. Gigascience 11.

37. Purushothaman P, Thakker S, Verma SC. 2015. Transcriptome Analysis of Kaposi’s Sarcoma-Associated Herpesvirus during De Novo Primary Infection of Human B and Endothelial Cells. J Virol 89:3093–3111.

38. Strahan R, Uppal T, Verma S. 2016. Next-Generation Sequencing in the Understanding of Kaposi’s Sarcoma-Associated Herpesvirus (KSHV) Biology. Viruses 8:92.

39. McClure L V., Kincaid RP, Burke JM, Grundhoff A, Sullivan CS. 2013. Comprehensive Mapping and Analysis of Kaposi’s Sarcoma-Associated Herpesvirus 3′ UTRs Identify Differential Posttranscriptional Control of Gene Expression in Lytic versus Latent Infection. J Virol 87:12838– 12849.

40. Bai Z, Huang Y, Li W, Zhu Y, Jung JU, Lu C, Gao S-J. 2014. Genomewide Mapping and Screening of Kaposi’s Sarcoma-Associated Herpesvirus (KSHV) 3′ Untranslated Regions Identify Bicistronic and Polycistronic Viral Transcripts as Frequent Targets of KSHV MicroRNAs. J Virol 88:377–392.

41. Arias C, Weisburd B, Stern-Ginossar N, Mercier A, Madrid AS, Bellare P, Holdorf M, Weissman JS, Ganem D. 2014. KSHV 2.0: A Comprehensive Annotation of the Kaposi’s Sarcoma-Associated Herpesvirus Genome Using Next-Generation Sequencing Reveals Novel Genomic and Functional Features. PLoS Pathog 10:e1003847.

42. Batut P, Gingeras TR. 2013. RAMPAGE: Promoter Activity Profiling by Paired-End Sequencing of 5′-Complete cDNAs. Curr Protoc Mol Biol 104.

43. Ye X, Zhaoid Y, Karijolich J. 2019. The landscape of transcription initiation across latent and lytic KSHV genomes. PLoS Pathog 15:1–26.

44. Zhao Y, Ye X, Shehata M, Dunker W, Xie Z, Karijolich J. 2020. The RNA quality control pathway nonsense-mediated mRNA decay targets cellular and viral RNAs to restrict KSHV. Nat Commun 11:3345.

45. Majerciak V, Alvarado-Hernandez B, Lobanov A, Cam M, Zheng Z-M. 2022. Genome-wide regulation of KSHV RNA splicing by viral RNA-binding protein ORF57. PLoS Pathog 18:e1010311.

46. Balázs Z, Tombácz D, Csabai Z, Moldován N, Snyder M, Boldogkoi Z. 2019. Template-switching artifacts resemble alternative polyadenylation. BMC Genomics 10.1186/s12864-019-6199-7.

47. Wyman D, Balderrama-Gutierrez G, Reese F, Jiang S, Rahmanian S, Forner S, Matheos D, Zeng W, Williams B, Trout D, England W, Chu S-H, Spitale RC, Tenner AJ, Wold BJ, Mortazavi A. 2020. A technology-agnostic long-read analysis pipeline for transcriptome discovery and quantification. bioRxiv 672931.

48. Majerciak V, Ni T, Yang W, Meng B, Zhu J, Zheng Z-M. 2013. A Viral Genome Landscape of RNA Polyadenylation from KSHV Latent to Lytic Infection. PLoS Pathog 9:e1003749.

49. Frith MC, Valen E, Krogh A, Hayashizaki Y, Carninci P, Sandelin A. 2008. A code for transcription initiation in mammalian genomes. Genome Res 18:1–12.

50. Ni T, Corcoran DL, Rach EA, Song S, Spana EP, Gao Y, Ohler U, Zhu J. 2010. A paired-end sequencing strategy to map the complex landscape of transcription initiation. Nat Methods 7:521– 527.

51. Calvo-Roitberg E, Daniels RF, Pai AA. 2023. Challenges in identifying mRNA transcript starts and ends from long-read sequencing data. bioRxiv, Prepr Serv Biol 10.1101/2023.07.26.550536.

52. Tombácz D, Prazsák I, Csabai Z, Moldován N, Dénes B, Snyder M, Boldogkői Z. 2020. Long-read assays shed new light on the transcriptome complexity of a viral pathogen. Sci Rep 10.1038/s41598-020-70794-5.

53. Unal A, Pray TR, Lagunoff M, Pennington MW, Ganem D, Craik CS. 1997. The protease and the assembly protein of Kaposi’s sarcoma-associated herpesvirus (human herpesvirus 8). J Virol 71:7030–7038.

54. Dresang LR, Teuton JR, Feng H, Jacobs JM, Camp DG, Purvine SO, Gritsenko MA, Li Z, Smith RD, Sugden B, Moore PS, Chang Y. 2011. Coupled transcriptome and proteome analysis of human lymphotropic tumor viruses: Insights on the detection and discovery of viral genes. BMC Genomics 10.1186/1471-2164-12-625.

55. Boldogkői Z, Moldován N, Balázs Z, Snyder M, Tombácz D. 2019. Long-Read Sequencing – A Powerful Tool in Viral Transcriptome Research. Trends Microbiol 10.1016/j.tim.2019.01.010.

56. Taylor JL, Bennett HN, Snyder BA, Moore PS, Chang Y. 2005. Transcriptional Analysis of Latent and Inducible Kaposi’s Sarcoma-Associated Herpesvirus Transcripts in the K4 to K7 Region. J Virol 79:15099–15106.

57. Bruce AG, Barcy S, DiMaio T, Gan E, Garrigues HJ, Lagunoff M, Rose TM. 2017. Quantitative Analysis of the KSHV Transcriptome Following Primary Infection of Blood and Lymphatic Endothelial Cells. Pathog (Basel, Switzerland) 6.

58. Landis JT, Tuck R, Pan Y, Mosso CN, Eason AB, Moorad R, Marron JS, Dittmer DP. 2022. Evidence for Multiple Subpopulations of Herpesvirus-Latently Infected Cells. MBio 13.

59. Depledge DP, Breuer J. 2021. Varicella-Zoster Virus—Genetics, Molecular Evolution and Recombination, p. 1–23. In 2021: pp. 1–23. 10.1007/82_2021_238.

60. Tombácz D, Moldován N, Balázs Z, Gulyás G, Csabai Z, Boldogkői M, Snyder M, Boldogkői Z. 2019. Multiple Long-Read Sequencing Survey of Herpes Simplex Virus Dynamic Transcriptome. Front Genet 10.3389/fgene.2019.00834.

61. Moldován N, Torma G, Gulyás G, Hornyák Á, Zádori Z, Jefferson VA, Csabai Z, Boldogkői M, Tombácz D, Meyer F, Boldogkői Z. 2020. Time-course profiling of bovine alphaherpesvirus 1.1 transcriptome using multiplatform sequencing. Sci Rep 10.1038/s41598-020-77520-1.

62. Torma G, Tombácz D, Csabai Z, Göbhardter D, Deim Z, Snyder M, Boldogkői Z. 2021. An Integrated Sequencing Approach for Updating the Pseudorabies Virus Transcriptome. Pathogens 10:242.

63. Fülöp Á, Torma G, Moldován N, Szenthe K, Bánáti F, Almsarrhad IAA, Csabai Z, Tombácz D, Minárovits J, Boldogkői Z. 2022. Integrative profiling of Epstein–Barr virus transcriptome using a multiplatform approach. Virol J 19:1–17.

64. Boldogkői Z, Balázs Z, Moldován N, Prazsák I, Tombácz D. 2019. Novel classes of replication-associated transcripts discovered in viruses. RNA Biol 10.1080/15476286.2018.1564468.

65. Tombácz D, Balázs Z, Csabai Z, Moldován N, Szűcs A, Sharon D, Snyder M, Boldogköi Z. 2017. Characterization of the Dynamic Transcriptome of a Herpesvirus with Long-read Single Molecule Real-Time Sequencing. Sci Rep 10.1038/srep43751.

66. Prusty BK, Whisnant AW. 2020. Revisiting the genomes of herpesviruses. Elife 9.

67. Brulois K, Toth Z, Wong L-Y, Feng P, Gao S-J, Ensser A, Jung JU. 2014. Kaposi’s Sarcoma-Associated Herpesvirus K3 and K5 Ubiquitin E3 Ligases Have Stage-Specific Immune Evasion Roles during Lytic Replication. J Virol 88:9335–9349.

68. Zhu FX, Cusano T, Yuan Y. 1999. Identification of the Immediate-Early Transcripts of Kaposi’s Sarcoma-Associated Herpesvirus. J Virol 73:5556–5567.

69. Wang Y, Li H, Chan MY, Zhu FX, Lukac DM, Yuan Y. 2004. Kaposi’s sarcoma-associated herpesvirus ori-Lyt-dependent DNA replication: cis-acting requirements for replication and ori-Lyt-associated RNA transcription. J Virol 78:8615–29.

70. Campbell M, Izumiya Y. 2020. PAN RNA: transcriptional exhaust from a viral engine. J Biomed Sci 27:41.

71. Schifano JM, Corcoran K, Kelkar H, Dittmer DP. 2017. Expression of the Antisense-to-Latency Transcript Long Noncoding RNA in Kaposi’s Sarcoma-Associated Herpesvirus. J Virol 91.

72. Xu Y, Ganem D. 2010. Making Sense of Antisense: Seemingly Noncoding RNAs Antisense to the Master Regulator of Kaposi’s Sarcoma-Associated Herpesvirus Lytic Replication Do Not Regulate That Transcript but Serve as mRNAs Encoding Small Peptides. J Virol 84:5465–5475.

73. Savoret J, Mesnard J-M, Gross A, Chazal N. 2021. Antisense Transcripts and Antisense Protein: A New Perspective on Human Immunodeficiency Virus Type 1. Front Microbiol 11.

74. Whisnant AW, Jürges CS, Hennig T, Wyler E, Prusty B, Rutkowski AJ, L’hernault A, Djakovic L, Göbel M, Döring K, Menegatti J, Antrobus R, Matheson NJ, Künzig FWH, Mastrobuoni G, Bielow C, Kempa S, Liang C, Dandekar T, Zimmer R, Landthaler M, Grässer F, Lehner PJ, Friedel CC, Erhard F, Dölken L. 2020. Integrative functional genomics decodes herpes simplex virus 1. Nat Commun 11:1–14.

75. Root-Bernstein RS, Holsworth DD. 1998. Antisense Peptides: A Critical Mini-Review. J Theor Biol 190:107–119.

76. Tagawa T, Serquiña A, Kook I, Ziegelbauer J. 2021. Viral non-coding RNAs: Stealth strategies in the tug-of-war between humans and herpesviruses. Semin Cell Dev Biol 111:135–147.

77. Tagawa T, Oh D, Dremel S, Mahesh G, Koparde VN, Duncan G, Andresson T, Ziegelbauer JM. 2023. A virus-induced circular RNA maintains latent infection of Kaposi’s sarcoma herpesvirus. Proc Natl Acad Sci 120.

78. Tagawa T, Oh D, Santos J, Dremel S, Mahesh G, Uldrick TS, Yarchoan R, Kopardé VN, Ziegelbauer JM. 2021. Characterizing Expression and Regulation of Gamma-Herpesviral Circular RNAs. Front Microbiol 12.

79. Martin-Baniandres P, Lan W-H, Board S, Romero-Ruiz M, Garcia-Manyes S, Qing Y, Bayley H. 2023. Enzyme-less nanopore detection of post-translational modifications within long polypeptides. Nat Nanotechnol 10.1038/s41565-023-01462-8.

80. Yu L, Kang X, Li F, Mehrafrooz B, Makhamreh A, Fallahi A, Foster JC, Aksimentiev A, Chen M, Wanunu M. 2023. Unidirectional single-file transport of full-length proteins through a nanopore. Nat Biotechnol 41:1130–1139.

81. Papp B, Motlagh N, Smindak RJ, Jin Jang S, Sharma A, Alonso JD, Toth Z. 2019. Genome-Wide Identification of Direct RTA Targets Reveals Key Host Factors for Kaposi’s Sarcoma-Associated Herpesvirus Lytic Reactivation. J Virol 93:1–22.

82. Toth Z, Brulois K, Lee HR, Izumiya Y, Tepper C, Kung HJ, Jung JU. 2013. Biphasic Euchromatin-to-Heterochromatin Transition on the KSHV Genome Following De Novo Infection. PLoS Pathog 9:1–14.

